# Environmental antibiotic resistome: selection of HAE-1 efflux pumps in early stages of plant root development

**DOI:** 10.1101/2024.09.04.611341

**Authors:** Olwen Simenel, Sylvie Nazaret, Josselin Bodilis

## Abstract

Natural environments are known to be reservoirs for antibiotic resistance genes (ARGs) and human pathogens. Thus, the One Health approach is necessary to fully understand their origin, spread and evolution. Among environments, the rhizosphere – i.e. the volume of soil in contact with plant roots - is of particular interest as it is both a hotspot of bacterial biomass and activity, and ARGs. In this study we investigated the role of the rhizosphere in the selection of antibiotic resistance in its associated bacterial community. We compared the abundance and expression levels of ARGs in seven metagenomes obtained after isotope labeling (DNA-SIP) and eighteen metatranscriptomes of soil and rhizosphere of *Avena fatua* from six to nine weeks old. ARGs were identified using the CARD database and an in-house HAE-1 (Hydrophobe Amphiphile Efflux-1) efflux pumps database. Our results showed that genes encoding the HAE-1 efflux pumps are enriched in the rhizosphere of six- and nine-week-old plants among the bacteria metabolizing the root exudates, and significantly overexpressed in the rhizosphere of nine-week-old plants. Additionnally, the diversity of HAE-1 pumps expressed in the rhizosphere was observed to be considerable, encompassing the full range of known diversity of these pumps in Gram-negative bacteria. We demonstrated that the environmental conditions in the rhizosphere directly selects for the HAE-1 pumps that is a major multidrug resistance factor in Gram-negative human pathogens. Studying the resistome of the rhizosphere is thus important to understand the emergence of multidrug resistance in human opportunistic pathogens.

## Introduction

Natural environments are known to be reservoirs of human pathogens which can be transmitted to people through diverse routes [1-3]. Of greater concern is the development and spread of bacterial antibiotic resistances in hospital and also in natural environments [4]. As a consequence of common and excessive antibiotic use, numerous multi-resistant enteric bacteria are released into the environment via faecal discharges originating from humans or animals [5,6]. Additionally, antibiotic resistance genes existed long before the use of antibiotics by human [7,8]. Thus, adopting a One Health approach, i.e. in a global environmental context, is essential to investigate the emergence of pathogenic bacteria and their antibiotic resistance mechanisms.

The rhizosphere, i.e. the volume of soil in contact with the plant roots, and under their physical and chemical influence, is nutrient-rich due to root exudates, which can represent a significant proportion of the plant’s photosynthetized carbon [9,10]. This abundance of nutrients favors a higher bacterial density and activity than in bulk (non-rhizospheric) soil [11]. The rhizosphere microbiome is very important for crop health, as its microorganisms interact with plants in numerous ways. Plant Growth-Promoting Rhizobacteria (PGPR) and fungi compete against plant pathogens [12] and help the plant to assimilate nitrogen and phosphorus [13-15].

The rhizosphere and bulk soils have been described as reservoirs of antibiotic resistance genes (ARG).^16,17^ Most studies on the subject focus on agricultural soils, particularly those that have been amended with manure, wastewater or struvite [18,19]. However, more recent studies have demonstrated that ARGs are enriched in the rhizosphere even in absence of treatments [20-22].

At least three reasons, not necessarily mutually exclusive, could explain the enrichment of ARGs in the rhizosphere. First, ARGs could be indirectly enriched in the rhizosphere because plants select specific bacteria belonging to genera or species well known for their resistance to antibiotics, such as *Pseudomonas spp*. or *Stenotrophomonas maltophilia* [22-26]. Second, the high microbial density in the rhizosphere generates competition between microorganisms, including antibiotic-producing organisms such as *Streptomycetes* and fungi [27,28], which could directly select ARGs in bacteria sharing their ecological niche. A third explanation is that some ARGs may be directly selected due to their broader physiological functions beyond their role in antibiotic resistance. These functions could be important for the adaptation of the bacteria to its environment [29-31].

In order to determine whether ARGs play a significant role and are therefore directly selected in the rhizosphere, it would be necessary to demonstrate firstly that they are carried by bacteria active in the rhizosphere and secondly that they are expressed, or even over-expressed, in comparison with the adjacent bulk soil.

To address this significant issue, we conducted a reanalysis of data from two studies based on the same model plant, *Avena fatua*, a common wild grass found in temperate regions on all continents, at the same growth stage, namely between six and nine weeks.

The first study involved Stable Isotope Probing (SIP) and metagenomic analyses, in which *A. fatua* is labelled with ^13^CO_2_ [32]. This study was selected for observation of the occurrence of ARGs in the active bacteria that had consumed the ^13^C photosynthetized from the plant.

The second study involved metatranscriptomic analyses on *A. fatua* grown in soil microcosms [33]. These data were re-analyzed to evaluate whether ARGs, and especially the HAE-1 efflux pumps, are (over)expressed in the rhizosphere.

## Methods

### Analysis of metagenomes and metatranscriptomes

We reanalyzed data from the study by Starr and colleagues 2021 [32]. The authors used the common annual grass *Avena fatua* as a model species. The plants were grown for six or nine weeks in microcosms within a growth chamber with a ^13^CO_2_-containing atmosphere. A single plant was utilized for each duration, with no replicates. The DNA from paired samples of rhizosphere and bulk soil were then extracted and separated by ultracentrifugation to obtain different fractions of DNA marked: fractions enriched in ^13^C, and (native) ^12^C fractions. The metagenomes from the ^12^C and ^13^C fractions of soil and rhizosphere, were reanalyzed, resulting in a total of seven metagenomes.

We also reanalyzed data from the study by Nuccio and colleagues 2020 [33]. *Avena fatua* was initially grown for six weeks in two-compartment microcosms with a mesh bag of soil serving as a bulk soil control. Then the wall between the two compartments was opened to allow the growth of roots into the second compartment. The growth of new roots was monitored for three weeks, with sampling of rhizosphere and bulk soil at three, six, 12 and 22 days after the opening of the second compartment. We analyzed the samples from the time points 1, 2 and 4, corresponding to three-, six-, and 22-days of experimentation, i.e. between six and nine weeks of plant growth, for a total of 18 samples.

### Treatment of the fastq files

The raw metatranscriptomic reads were downloaded from the Joint Genome Institute database, and the raw SIP-informed metagenomes were downloaded from the NCBI Sequence Read Archive using SRA-Toolkit, with forward and reverse reads of each sample in two separate fastq files. For each sample, forward and reverse reads were merged using the bbmerge.sh script from the BBTools suite [34], with the minavgquality option set to 20 to merge only reads with an average quality score of at least 20. Reads shorter than 210 base pairs were filtered using Mothur [35].

### Detection of *rpoB* genes of opportunistic pathogens

To assess the abundance and activity of opportunistic human pathogens in the SIP-informed metagenomes and metatranscriptomes, we quantified the *rpoB* genes, a single-copy housekeeping gene encoding the beta subunit of the RNA polymerase. Thus, the metatranscriptomes from Nuccio *et al*. 2020 were sequenced from 16S rRNA-depleted samples, which preclude the reliable use of the 16S rRNA for the detection of transcripts from potential human pathogens.

Two *rpo*B databases were created. The first database contains 506 protein sequences of the beta subunit of the RNA-polymerase, encompassing all phyla of Archaea, Gram-positive and Gram-negative bacteria. The second database, was constructed with 64 bacterial species that have been documented in the literature being associated with human infections and as having been isolated from rhizosphere (S1Table). The database was constructed using the nucleotides sequences of the *rpo*B gene of the reference strain of each of these species.

Subsequently, each metagenome and metatranscriptome was screened against the first database using BlastX with Blast standalone [36], with an e-value threshold of 10^−6^, and all hits with less than 70% sequence identity and 70 aminoacids of alignment length were discarded. The reads corresponding to the retained hits were extracted using the get.seqs function of Mothur, and screened against the second database by blastn with the same e-value threshold, and the hits with less than 95% sequence identity were discarded.

### Screening for antibiotic resistance genes and transcripts

To quantify ARGs in the metagenomes and metatranscriptomes, we mapped the filtered reads against the Comprehensive Antibiotic Resistance Database (CARD) using standalone BlastX [36]. Only hits with an e-value of less than 10^−6^, an identity percentage of at least 70% and an alignment length of at least 70 amino acids ere retained. The abundance of ARGs in hits per million reads was then calculated.

### Screening for HAE-1 efflux pumps

To identify HAE-1 efflux pumps with more precision, we mapped the metagenomes and metatranscriptomes using BlastX against an in-house database of 6205 protein sequences of RND efflux transporters encompassing the HAE-1, NFE and HME families. The sequences of our database form nine phylogenetic clades, which we previously defined in a study [20]. In this study, we proposed to restrict the HAE-1 family to two phylogenetic clades (A and B), which represent 41.8% of all RND pumps and group most of the RND pumps involved in multidrug resistance. Only those hits with an e-value of less than 10^−6^, at least 70% of sequence identity and an alignment length of at least 70 aminoacids were retained.

### Phylogenetic analysis of HAE-1 efflux pumps

From our initial database of 6,205 protein sequences of RND efflux transporters, we extracted the 2,592 sequences corresponding to HAE-1 (clade A and B) [20]. The 2,592 whole protein sequences were aligned using Clustal Omega 1.2.1 with the default parameters. The poorly aligned positions were eliminated using Gblock 0.91b with the default settings, except parameters allowing all positions and conserved alignment blocks of at least 50 columns. A protein alignment of 622 columns was then retained for phylogenetic analyses. Reconstruct Maximum Likelihood phylogenetic trees were obtained by using IQTree 1.6.5, with a maximum parsimony starting tree. The best-fitted model (LG+F+R4) was selected based on Bayesian Information Criterion and 1,000 samples were used to determine ultrafast bootstraps.

### Statistical analysis

The statistical analyses were performed with R 4.2.2 [37] The relative abundances and expression levels of HAE-1 efflux pumps were compared using either the paired t-test, or the Wilcoxon test depending on whether the required conditions for applying the t-test were met. A difference was regarded as significant if the p-value was at most 0.05.

## Results

### Treatment of the metagenomes and metatranscriptomes

After merging the forward and reverse reads, and discarding the reads of low-quality or too short, we kept between 4,891,760 and 7,700,594 reads per metagenome (5,973,911 reads on average), and between 6,221,809 and 18,705,403 reads per metatranscriptome (11,252,761 reads on average).

### Presence and overexpression of ARGS in the rhizosphere

We detected ARGs in all rhizosphere metagenomes, both in the ^12^C and ^13^C fractions of DNA, i.e. corresponding to bacteria that had consumed the ^13^C photosynthetized from the plant (S2 Table). Interestingly, the ^12^C fractions of rhizosphere DNA showed a total abundance of 202.55 and 252.05 hits per million reads at six and nine weeks, respectively, while the ^13^C fractions of rhizosphere DNA showed 562.33 hits per million reads at six weeks and 662.06 at nine weeks (Table 1).

**Table 1.** relative abundance of the top 10 ARGs in bulk soil and rhizosphere metagenomes. The total HAE-1 refers to hits with the RND database

|  | t0 | 6 weeks |  |  | 9 weeks |  |  |
| --- | --- | --- | --- | --- | --- | --- | --- |
|  | bulk, <sup>12</sup>C | bulk, <sup>12</sup>C | rhizo, <sup>12</sup>C | rhizo, <sup>13</sup>C | bulk, <sup>12</sup>C | rhizo, <sup>12</sup>C | rhizo, <sup>13</sup>C |
| *muxB* | 30,63 | 31,84 | 21,42 | 44,34 | 37,19 | 31,32 | 53,49 |
| *mdtB* | 24,18 | 19,66 | 14,77 | 38,13 | 21,59 | 18,01 | 50,73 |
| *ceoB* | 10,80 | 8,18 | 6,82 | 54,18 | 8,34 | 7,47 | 59,53 |
| *mexF* | 21,60 | 17,22 | 12,01 | 26,74 | 16,69 | 19,80 | 33,47 |
| *adeF* | 12,41 | 9,22 | 6,98 | 49,52 | 9,07 | 10,71 | 49,00 |
| *mdtC* | 20,15 | 19,49 | 13,15 | 27,26 | 15,96 | 17,85 | 29,16 |
| *novA* | 10,96 | 19,14 | 9,09 | 7,94 | 12,15 | 11,52 | 13,80 |
| *vanRO* | 13,38 | 12,70 | 10,87 | 6,04 | 10,70 | 12,17 | 13,11 |
| *mexB* | 5,00 | 3,48 | 3,90 | 31,06 | 2,72 | 3,41 | 26,92 |
| *mexK* | 6,29 | 4,35 | 5,36 | 22,09 | 5,62 | 5,19 | 26,05 |
| Total ARGs (CARD database) | 275,68 | 268,27 | 202,55 | 562,33 | 263,41 | 252,05 | 662,06 |
| total HAE-1 | 852,52 | 752,62 | 637,83 | 1631,77 | 804,93 | 833,56 | 1715,11 |

Among the 10 most abundant ARGs detected, eight encode RND efflux pumps. These genes appeared to be the most enriched in the ^13^C rhizosphere fractions (S1 Fig).

In terms of ARGS expression, a BlastX search against the CARD database showed a total relative abundance of ARGs per metatranscriptome, ranging from 58.63 to 91.95 hits per million non-16S rRNA reads.

### Enrichment and overexpression of RND efflux pumps of the HAE-1 family in the rhizosphere

The RND efflux pumps are divided in eight families, three of which (HAE-1, HME and NFE) are closely related, and difficult to discriminate [20]. We also observed that the annotation using the CARD database is not precise enough to correctly identify the pumps belonging to these families. Given the central role of HAE-1 family in multidrug resistance in Gram-negative bacteria, we focused our analysis on this efflux pump family by using an in-house database covering the diversity of HAE-1 pumps of environmental bacteria [20]. We detected genes coding HAE-1 efflux pumps in all metagenomes of soil and rhizosphere. Interestingly, the relative abundance of HAE-1 genes in the ^12^C fractions of rhizospheric DNA was similar to that of soil, with 637.83 hits per million reads at six weeks and 833.56 hits per million reads at nine weeks, while in the ^13^C fractions of rhizospheric DNA the HAE-1 genes were approximately twice more abundant, with 1631.77 and 1715.11 hits per million reads at six and nine weeks, respectively (Table 1).

Transcripts of HAE-1 pumps were detected in all metatranscriptomes, in bulk soil and rhizosphere samples. Interestingly, the relative abundance of HAE-1 transcripts in the rhizosphere was higher than in bulk soil as early as three days of root growth, or six weeks, and this difference became significant at 22 days of root growth, corresponding to nine weeks of age, where HAE-1 transcripts represented 243.4 HAE-1 reads per million non-16S rRNA reads in the rhizosphere, vs 175 HAE-1 reads per million non-16S rRNA reads in the bulk soil (p=0.05, paired t test, see Figure 1).

**Fig 1.**
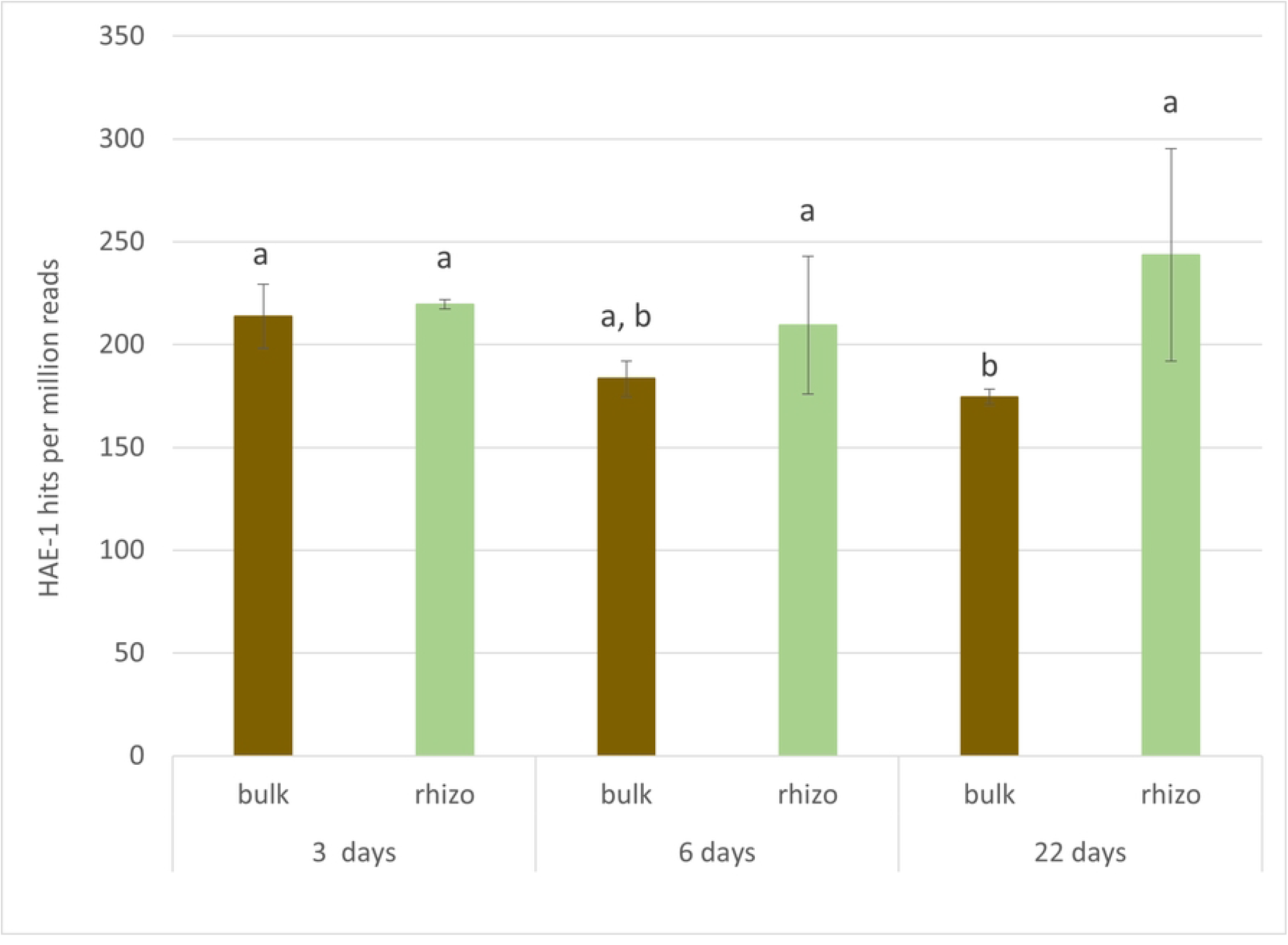
Relative abundances of HAE-1 reads in bulk soil and in rhizosphere. Brown : bulk soil, Green : rhizosphere

### Abundance and activity of opportunistic pathogens in soil and rhizosphere

We detected *rpoB* genes with at least 95% identity to opportunistic pathogens in all soil and rhizosphere metagenomes, including those derived from the ^13^C-enriched DNA and ^12^C DNA (S1 Table). In soil metagenomes (i.e. without the plant), the *rpoB* sequences from opportunistic pathogens represented 0.65% of all detected *rpo*B sequences at t0. However, in the metagenomes from the ^13^C fraction of rhizosphere DNA, corresponding to rhizosphere bacteria that did consume the root exudates, the relative abundance of *rpo*B sequences from opportunistic pathogens was higher, at 0.77% and 0.78% at six and nine weeks, respectively (S3 Table). Due to the absence of replicates, it is not possible to perform statistical analyses on these results. However, the data suggest that the root exudates select for opportunistic pathogenic bacteria.

We also detected *rpo*B transcripts of opportunistic pathogens in all soil and rhizosphere metatranscriptomes representing between 0.30% and 0.49% of all detected *rpo*B transcripts. No significant differences were detected between soil and rhizosphere metatranscriptomes at any time point (S3 Table).

### High diversity of HAE-1 pumps in the rhizosphere

Among the bacteria that have consumed the ^13^C-marked root exudates, we detected 934 HAE-1 transporter genes (out of 2,592 in our RND database), of which 758 (>80%) were also detected in the *A. fatua* rhizosphere metatranscriptomes. Overall, more than half of the known diversity of HAE-1 transporters (1,320 out of 2,592) is expressed and/or present in the genomes of bacteria metabolizing root exudates. Surprisingly, even though our RND database is based on HAE-1 transporters from phylogenetically and ecologically diverse species, the HAE-1s expressed in the rhizosphere are distributed throughout the phylogenetic tree (Fig 2). In addition, among our HAE-1 database, we have identified 19 pumps described in the literature as being involved in multidrug resistance in the clinical setting. These pumps are distributed throughout the phylogenetic tree in a similar manner, with more than half being present and/or expressed in the rhizosphere (Fig 2).

**Fig 2.**
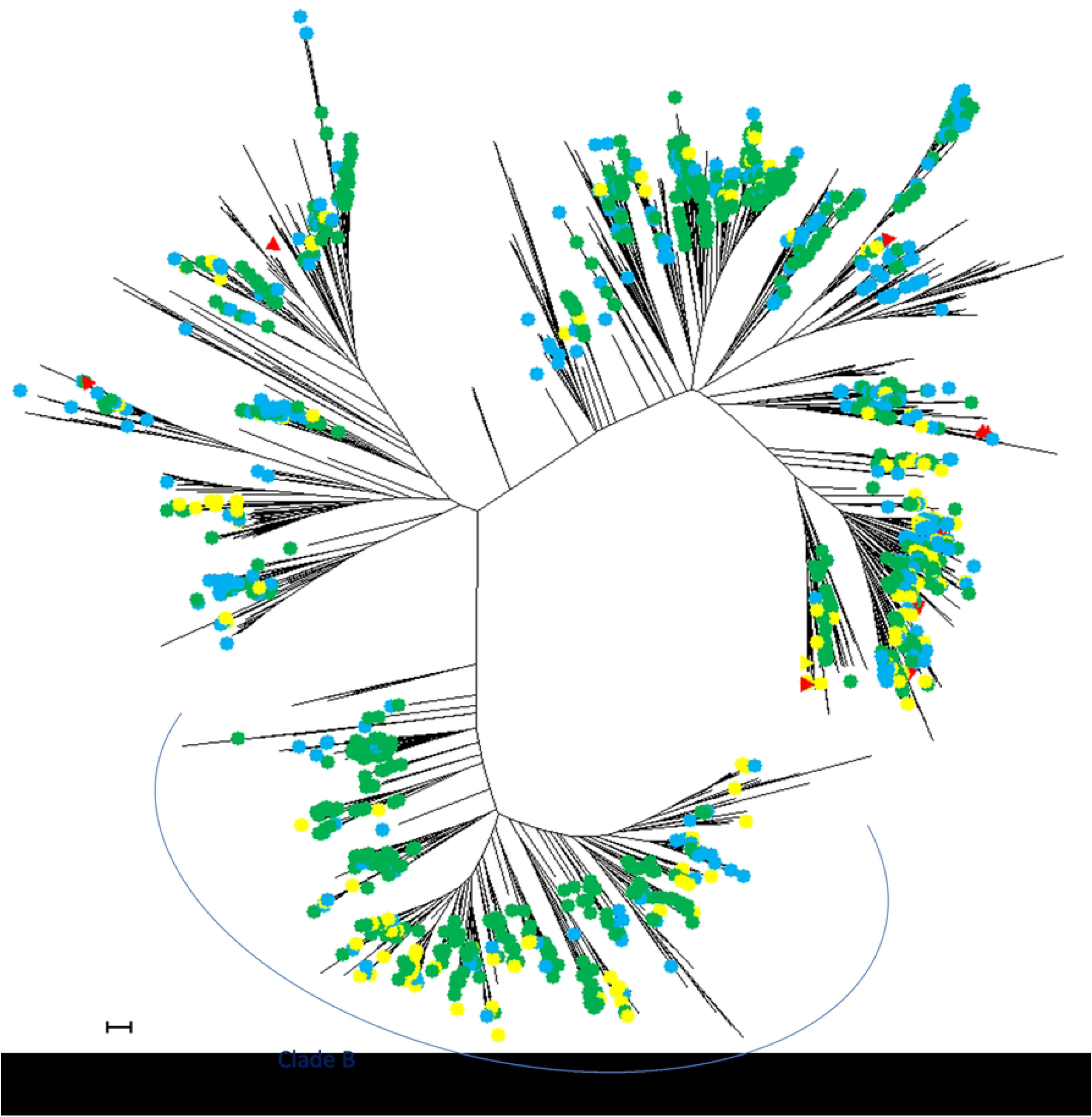
Phylogenetic tree from 2592 HAE-1 permeases. Sequences were obtained from 920 reference proteomes of Gram-negative bacteria (Uniprot, release 2015_09). Colored dots indicate HAE-1 pumps present in bacteria that have consumed the 13C-marked root exudates (in yellow), HAE-1 pumps expressed in the rhizosphere (in blue) or both present and expressed (in green). Pumps already described as being involved in multiresistance to antibiotics are indicated by a red delta. Nodes with bootstrap values below 80% were collapsed.

## Discussion

### Rhizosphere directly selects and induces HAE-1 efflux pumps

In our metagenomic analysis, we observed that HAE-1 efflux pumps are the dominant class of ARGs in soil and rhizosphere, and seem to be enriched in rhizosphere bacteria metabolizing root exudates. These results are consistent with previous studies that have demonstrated that plants can select ARGs in their rhizosphere [21,22,25,38,39]. Besides a fitness advantage provided by the ARGs, several factors can explain their enrichment in the rhizosphere, such as the fact that ARG-carrying bacteria can be selected for other traits enhancing their fitness in rhizosphere [22], in which case the selection of ARGs in coincidental. In this context, overexpression of HAE-1 pumps is noteworthy because their expression is tightly regulated due to its high energetic cost. Most HAE-1 pumps are silent and can be induced only in presence of specific inducers like MexXY-OprM in *P. aeruginosa*, that is induced by agents targeting the ribosome [40], or in case of a mutation inactivating their repressor such as in *nfxB* and *nfxC* mutants in *P. aeruginosa* [41,42]. Other HAE-1 pumps such as MexAB-OprM in *P. aeruginosa* or SmeDEF in *S. maltophilia* are constitutively expressed but their expression is kept at a low level by several regulators [43-45]. Due to this tight regulation, an overexpression of HAE-1 pumps in the rhizosphere indicates they play an active role in this environment and their enrichment in rhizosphere bacteria is not coincidental.

Two hypotheses may explain the selection of HAE-1 pumps in the rhizosphere. Firstly, the HAE-1 efflux pumps may participate in the competition between microorganisms in the nutrient-rich environment of the rhizosphere. Secondly, they may provide resistance to the potentially toxic secondary plant metabolites present in root exudates.

### HAE-1 efflux pumps protect bacteria from toxic plant metabolites

Due to the high nutrient availability and the density of active microorganisms in the rhizosphere, bacteria in this environment are subjected to a high level of competition [11]. As most known natural antibiotics are produced by actinomycetes [27], that are abundant in telluric environments, HAE-1 efflux pumps could be a protection for Gram-negative bacteria against actinomycete competitors. This hypothesis is supported by the fact that most functionally characterized HAE-1 pumps extrude antibiotics, even in bacteria that are not pathogenic to humans and thus not exposed to high concentrations of antibiotics during infection such as environmental *Pseudomonas putida* and *P. stutzeri*, or *Bradyrhizobium japonicum* [46-48]. However, the concentrations of antibiotics in the rhizosphere may not be sufficient to exert an antibacterial effect, as there is a lack of available literature on the concentrations of antibiotics produced in the rhizosphere by micro-organisms. Furthermore, antibiotic contamination rarely reaches inhibitory concentrations [49], which suggests that antibiotics are unlikely to induce the expression of costly efflux pumps.

In addition to competition from other microorganisms, rhizosphere bacteria are subjected to the physico-chemical stresses generated from the plant roots. The roots can generate transient acid stress [50], and they excrete numerous secondary metabolites with diverse functions, some of which are defense molecules with antibacterial activities [51]. These stressors are the most likely cause of the overexpression and selection of HAE-1 efflux pumps in the rhizosphere, given that several HAE-1 efflux pumps are induced by stresses such as oxidative stress, acid stress or membrane damage, and can help mitigate these stresses [52-56]. Other HAE-1 pumps have been demonstrated to extrude plant secondary metabolites such as flavonoids. In some cases, the inactivation of the HAE-1 pump have been shown to decrease the resistance of the bacteria to flavonoids that are not inhibitory to the wild-type strain [57], or to decrease its fitness in the rhizosphere [57,58] or in the plant host during infection [59-61].

Plant secondary metabolites are highly diverse, and their concentration in root exudates varies depending on a number of factors including the plant species, its health status, and the presence of biotic or abiotic stresses. One such factor is the plant age, with the proportion of secondary metabolites in the exudates increasing with age [62-64]. This is observed in *Avena barbata*, a species related to *A. fatua*, where the exudation of secondary metabolites reaches a maximum at nine weeks of age [64]. This could explain the observed overexpression of HAE-1 pumps at this time point. As some molecules present in the exudates of *A. barbata* and other plant species, like salicylic acid, have been demonstrated to induce the expression of pumps in bacteria [65], it is plausible that the high diversity of plant metabolites to which rhizosphere bacteria might be exposed could be related to the higher number of HAE-1 pumps with different substrate ranges and regulation mechanisms in their genomes compared to bacteria from bulk soil, which are exposed to less diverse molecules.

### The rhizosphere provides a pre-adaptation of bacteria to antibiotics

Our phylogenetic analysis shows that the HAE-1 pumps involved in rhizosphere colonization are distribute across the phylogenetic tree of HAE-1 pumps, rather than forming a distinct cluster related to plant colonization. This suggest that most, if not all, HAE-1 pumps involved in multidrug resistance in clinical settings are related to a rhizosphere-related HAE-1 pump (Fig 2). Additionally, 22 opportunistic pathogens of our list (S1 Table) are represented in our RND database. Of these, 19 have at least one HAE-1 pump expressed in the rhizosphere, which confirms the dual role of HAE-1 pumps in infection and rhizosphere colonization.

The results of our phylogenetic, metagenomic and metatranscriptomic analyses suggest that the rhizosphere provides a pre-adaptation of the rhizosphere bacteria to antibiotics, by selecting a high number of HAE-1 pumps in their genomes.

In Gram-negative opportunistic pathogens, we argue that there is a physiological equilibrium between to distinct expression patterns : (i) the expression of a single generalist HAE-1 pump at a high level, and (ii) expression of multiple HAE-1 pumps at a low level or only in specific conditions. Most environmental strains and clinical isolates of these bacteria fall into the second configuration, while multidrug resistant clinical isolates fall into the first one, i.e. one generalist HAE-1 pump that is constitutively expressed at a high level due to an inactivation of its repressor. Due to the wide substrate spectrum of the overexpressed pump, this configuration causes loss of virulence or fitness [66] through metabolite leaks, but it provides a high level of resistance against antibiotics, which are the primary selective pressure during infection and treatment. Understanding this evolutionary event – the point where the advantages of the HAE-1 pump deregulation compensate the fitness costs – is important to improve antibiotic treatment courses to prevent the emergence of multidrug resistant mutants, especially in patient with cystic fibrosis, as they often have chronic infections by Gram-negative pathogens such as *P. aeruginosa* and repeated antibiotic treatments. It is also important to understand the mechanisms of the pre-adaptation of rhizosphere bacteria to antibiotics in order to limit the emergence and transmission of HAE-1-rich opportunistic pathogenic bacteria from crops to humans.

### Concluding remarks

This study demonstrates that environmental conditions in the rhizosphere directly select for a high number of HAE-1 efflux pumps in rhizosphere bacteria. These efflux pumps play a role in the colonization of the rhizosphere, but they also represent a significant antibiotic resistance factor in human pathogens such as *P. aeruginosa*. Studying the resistome of the rhizosphere is thus important to understand how plants build up their root microbiota and for developing alternatives to chemical inputs in agriculture, for understanding and reducing the emergence of multidrug resistance in human opportunistic pathogens.

## Supporting Information

**S1 Fig. Relative abundances of ARGs in** ^**12**^**C and** ^**13**^**C fractions of bulk soil and rhizospheric DNA**

**S2 Fig. Relative abundances of HAE-1 transporters in** ^**12**^**C and** ^**13**^**C fractions of bulk soil and rhizospheric DNA**

Bulk : bulk soil, rhizo : rhizosphere

**S1 Table. List of opportunistic human pathogens present in the rhizosphere and references**

**S2 Table. Relative abundance of ARGs in** ^**12**^**C and** ^**13**^**C fractions of bulk soil and rhizospheric DNA**

Values are in hits per million reads

**S3 Table. Relative abundance of rpoB genes from opportunistic pathogenes** values are in percentage of total rpoB sequences detected, values for metatranscriptomes are averages of three samples

## References

1. Baumgardner DJ. Soil-Related Bacterial and Fungal Infections. The Journal of the American Board of Family Medicine. 1 sept 2012;25(5):734–44.

2. Forstinus N, Ikechukwu N, Emenike M, Christiana A. Water and Waterborne Diseases: A Review. IJTDH. 10 janv 2016;12(4):1–14.

3. Sobiczewski P, Iakimova ET. Plant and Human Pathogenic Bacteria Exchanging their Primary Host Environments. Journal of Horticultural Research. 1 juin 2022;30(1):11–30.

4. Larsson DGJ, Flach CF. Antibiotic resistance in the environment. Nat Rev Microbiol. mai 2022;20(5):257–69.

5. Marutescu LG, Popa M, Gheorghe-Barbu I, Barbu IC, Rodríguez-Molina D, Berglund F, et al. Wastewater treatment plants, an “escape gate” for ESCAPE pathogens. Front Microbiol [Internet]. 2023;14.

6. Zalewska M, Błażejewska A, Czapko A, Popowska M. Antibiotics and Antibiotic Resistance Genes in Animal Manure – Consequences of Its Application in Agriculture. Front Microbiol. 2021

7. D’Costa VM, King CE, Kalan L, Morar M, Sung WWL, Schwarz C, et al. Antibiotic resistance is ancient. Nature. sept 2011;477(7365):457–61.

8. Van Goethem MW, Pierneef R, Bezuidt OKI, Van De Peer Y, Cowan DA, Makhalanyane TP. A reservoir of ‘historical’ antibiotic resistance genes in remote pristine Antarctic soils. Microbiome. 23 févr 2018;6(1):40.

9. Bais HP, Weir TL, Perry LG, Gilroy S, Vivanco JM. The role of root exudates in rhizosphere interactions with plants and other organisms. Annu Rev Plant Biol. 1 juin 2006;57(1):233–66.

10. Doornbos RF, van Loon LC, Bakker Pahm. Impact of root exudates and plant defense signaling on bacterial communities in the rhizosphere. A review. Agron Sustain Dev. 1 janv 2012;32(1):227–43.

11. Foster RC. Microenvironments of soil microorganisms. Biol Fert Soils. 1 mai 1988;6(3):189–203.

12. Lugtenberg B, Kamilova F. Plant-Growth-Promoting Rhizobacteria. Annu Rev Microbiol. 1 oct 2009;63(1):541–56.

13. Mendes R, Garbeva P, Raaijmakers JM. The rhizosphere microbiome: significance of plant beneficial, plant pathogenic, and human pathogenic microorganisms. FEMS Microbiology Reviews. 1 sept 2013;37(5):634–63.

14. Pereira SIA, Castro PML. Phosphate-solubilizing rhizobacteria enhance Zea mays growth in agricultural P-deficient soils. Ecological Engineering. 1 déc 2014;73:526–35.

15. Vega NWO. A review on beneficial effects of rhizosphere bacteria on soil nutrient availability and plant nutrient uptake. 2007

16. Heuer H, Krögerrecklenfort E, Wellington EMH, Egan S, van Elsas JD, van Overbeek L, et al. Gentamicin resistance genes in environmental bacteria: prevalence and transfer. FEMS Microbiology Ecology. 1 nov 2002;42(2):289–302.

17. Qian X, Gunturu S, Guo J, Chai B, Cole JR, Gu J, et al. Metagenomic analysis reveals the shared and distinct features of the soil resistome across tundra, temperate prairie, and tropical ecosystems. Microbiome. 14 mai 2021;9(1):108.

18. Chen QL, An XL, Zhu YG, Su JQ, Gillings MR, Ye ZL, et al. Application of Struvite Alters the Antibiotic Resistome in Soil, Rhizosphere, and Phyllosphere. Environ Sci Technol. 18 juill 2017;51(14):8149–57.

19. Urra J, Alkorta I, Mijangos I, Epelde L, Garbisu C. Application of sewage sludge to agricultural soil increases the abundance of antibiotic resistance genes without altering the composition of prokaryotic communities. Science of The Total Environment. 10 janv 2019;647:1410–20.

20. Bodilis J, Simenel O, Michalet S, Brothier E, Meyer T, Favre-Bonté S, et al. HME, NFE, and HAE-1 efflux pumps in Gram-negative bacteria: a comprehensive phylogenetic and ecological approach. ISME Communications. 1 janv 2024;4(1):ycad018.

21. Shami AY, Abulfaraj AA, Refai MY, Barqawi AA, Binothman N, Tashkandi MA, et al. Abundant antibiotic resistance genes in rhizobiome of the human edible Moringa oleifera medicinal plant. Front Microbiol. 15 sept 2022;13:990169.

22. Yu Y, Zhang Q, Zhang Z, Zhou S, Jin M, Zhu D, et al. Plants select antibiotic resistome in rhizosphere in early stage. Science of The Total Environment. 1 févr 2023;858:159847.

23. Youenou B, Favre-Bonté S, Bodilis J, Brothier E, Dubost A, Muller D, et al. Comparative Genomics of Environmental and Clinical Stenotrophomonas maltophilia Strains with Different Antibiotic Resistance Profiles. Genome Biology and Evolution. 1 sept 2015;7(9):2484–505.

24. Berendsen RL, Vismans G, Yu K, Song Y, de Jonge R, Burgman WP, et al. Disease-induced assemblage of a plant-beneficial bacterial consortium. ISME J. juin 2018;12(6):1496–507.

25. Liu B, Zhang D, Pan X. Nodules of wild legumes as unique natural hotspots of antibiotic resistance genes. Science of The Total Environment. 15 sept 2022;839:156036.

26. Sasse J, Martinoia E, Northen T. Feed Your Friends: Do Plant Exudates Shape the Root Microbiome? Trends in Plant Science. 1 janv 2018;23(1):25–41.

27. Williams ST, Vickers JC. The ecology of antibiotic production. Microb Ecol. mars 1986;12(1):43–52.

28. van der Meij A, Worsley SF, Hutchings MI, van Wezel GP. Chemical ecology of antibiotic production by actinomycetes. FEMS Microbiology Reviews. 1 mai 2017;41(3):392–416.

29. Alvarez-Ortega C, Olivares J, Martinez J. RND multidrug efflux pumps: what are they good for? Frontiers in Microbiology [Internet]. 2013 4.

30. Henderson PJF, Maher C, Elbourne LDH, Eijkelkamp BA, Paulsen IT, Hassan KA. Physiological Functions of Bacterial “Multidrug” Efflux Pumps. Chem Rev. 12 mai 2021;121(9):5417–78.

31. Martínez JL, Rojo F. Metabolic regulation of antibiotic resistance. FEMS Microbiology Reviews. 1 sept 2011;35(5):768–89.

32. Starr EP, Shi S, Blazewicz SJ, Koch BJ, Probst AJ, Hungate BA, et al. Stable-Isotope-Informed, Genome-Resolved Metagenomics Uncovers Potential Cross-Kingdom Interactions in Rhizosphere Soil. mSphere. sept 2021;6(5):10.1128/msphere.00085-21.

33. Nuccio EE, Starr E, Karaoz U, Brodie EL, Zhou J, Tringe SG, et al. Niche differentiation is spatially and temporally regulated in the rhizosphere. ISME J. avr 2020;14(4):999–1014.

34. Bushnell B, Rood J, Singer E. BBMerge – Accurate paired shotgun read merging via overlap. PLOS ONE. 26 oct 2017;12(10):e0185056.

35. Schloss P, Westcott S, Ryabin T, Hall J, Hartmann M, Hollister E, et al. Introducing mothur: Open-Source, Platform-Independent, Community-Supported Software for Describing and Comparing Microbial Communities [Internet]. 2009 [cité 20 mai 2022]. Disponible sur: https://journals.asm.org/doi/epdf/10.1128/AEM.01541-09

36. Camacho C, Coulouris G, Avagyan V, Ma N, Papadopoulos J, Bealer K, et al. BLAST+: architecture and applications. BMC Bioinformatics. 15 déc 2009;10(1):421.

37. R Core Team (2022). _R: A Language and Environment for Statistical Computing. R Foundation for Statistical Computing, Vienna, Austria. https://www.R-project.org.

38. Li X, Rui J, Xiong J, Li J, He Z, Zhou J, et al. Functional Potential of Soil Microbial Communities in the Maize Rhizosphere. PLOS ONE. 10 nov 2014;9(11):e112609.

39. Guo A, Pan C, Ma J, Bao Y. Linkage of antibiotic resistance genes, associated bacteria communities and metabolites in the wheat rhizosphere from chlorpyrifos-contaminated soil. Science of The Total Environment. 1 nov 2020;741:140457.

40. Jeannot K, Sobel ML, El Garch F, Poole K, Plésiat P. Induction of the MexXY Efflux Pump in Pseudomonas aeruginosa Is Dependent on Drug-Ribosome Interaction. Journal of Bacteriology. août 2005;187(15):5341–6.

41. Hirai K, Suzue S, Irikura T, Iyobe S, Mitsuhashi S. Mutations producing resistance to norfloxacin in Pseudomonas aeruginosa. Antimicrobial Agents and Chemotherapy. avr 1987;31(4):582–6.

42. Fukuda H, Hosaka M, Hirai K, Iyobe S. New norfloxacin resistance gene in Pseudomonas aeruginosa PAO. Antimicrobial Agents and Chemotherapy. sept 1990;34(9):1757–61.

43. Morita Y, Cao L, Gould VC, Avison MB, Poole K. nalD Encodes a Second Repressor of the mexAB-oprM Multidrug Efflux Operon of Pseudomonas aeruginosa. Journal of Bacteriology. 15 déc 2006;188(24):8649–54.

44. Sánchez P, Alonso A, Martinez JL. Cloning and Characterization of SmeT, a Repressor of the Stenotrophomonas maltophilia Multidrug Efflux Pump SmeDEF. Antimicrobial Agents and Chemotherapy. nov 2002;46(11):3386–93.

45. Tian ZX, Yi XX, Cho A, O’Gara F, Wang YP. CpxR Activates MexAB-OprM Efflux Pump Expression and Enhances Antibiotic Resistance in Both Laboratory and Clinical nalB-Type Isolates of Pseudomonas aeruginosa. PLOS Pathogens. 13 oct 2016;12(10):e1005932.

46. Alguel Y, Meng C, Terán W, Krell T, Ramos JL, Gallegos MT, et al. Crystal Structures of Multidrug Binding Protein TtgR in Complex with Antibiotics and Plant Antimicrobials. Journal of Molecular Biology. 8 juin 2007;369(3):829–40.

47. Jude F, Arpin C, Brachet-Castang C, Capdepuy M, Caumette P, Quentin C. TbtABM, a multidrug efflux pump associated with tributyltin resistance in Pseudomonas stutzeri. FEMS Microbiology Letters. 1 mars 2004;232(1):7–14.

48. Lindemann A, Koch M, Pessi G, Müller AJ, Balsiger S, Hennecke H, et al. Host-specific symbiotic requirement of BdeAB, a RegR-controlled RND-type efflux system in Bradyrhizobium japonicum. FEMS Microbiology Letters. 1 nov 2010;312(2):184–91.

49. Xie Y feng, Li XW, Wang JF, Christakos G, Hu MG, An LH, et al. Spatial estimation of antibiotic residues in surface soils in a typical intensive vegetable cultivation area in China. Science of The Total Environment. 15 juill 2012;430:126–31.

50. Weisskopf L, Fromin N, Tomasi N, Aragno M, Martinoia E. Secretion activity of white lupin’s cluster roots influences bacterial abundance, function and community structure. Plant Soil. 1 janv 2005;268(1):181–94.

51. Brigham LA, Michaels PJ, Flores HE. Cell-Specific Production and Antimicrobial Activity of Naphthoquinones in Roots of Lithospermum erythrorhizon1. Plant Physiology. 1 févr 1999;119(2):417–28.

52. Adebusuyi AA, Foght JM. An alternative physiological role for the EmhABC efflux pump in Pseudomonas fluorescens cLP6a. BMC Microbiol. 15 nov 2011;11(1):252.

53. Blanco P, Corona F, Sánchez MB, Martínez JL. Vitamin K3 Induces the Expression of the Stenotrophomonas maltophilia SmeVWX Multidrug Efflux Pump. Antimicrobial Agents and Chemotherapy. mai 2017;61(5):e02453–16.

54. Fetar H, Gilmour C, Klinoski R, Daigle DM, Dean CR, Poole K. mexEF-oprN Multidrug Efflux Operon of Pseudomonas aeruginosa: Regulation by the MexT Activator in Response to Nitrosative Stress and Chloramphenicol. Antimicrobial Agents and Chemotherapy. févr 2011;55(2):508–14.

55. Hou A ming, Yang D, Miao J, Shi D yang, Yin J, Yang Z wei, et al. Chlorine injury enhances antibiotic resistance in Pseudomonas aeruginosa through over expression of drug efflux pumps. Water Research. 1 juin 2019;156:366–71.

56. Huang YW, Liou RS, Lin YT, Huang HH, Yang TC. A Linkage between SmeIJK Efflux Pump, Cell Envelope Integrity, and σE-Mediated Envelope Stress Response in Stenotrophomonas maltophilia. PLOS ONE. 12 nov 2014;9(11):e111784.

57. García-León G, Hernández A, Hernando-Amado S, Alavi P, Berg G, Martínez JL. A Function of SmeDEF, the Major Quinolone Resistance Determinant of Stenotrophomonas maltophilia, Is the Colonization of Plant Roots. Applied and Environmental Microbiology. août 2014;80(15):4559–65.

58. Palumbo JD, Kado CI, Phillips DA. An Isoflavonoid-Inducible Efflux Pump in Agrobacterium tumefaciens Is Involved in Competitive Colonization of Roots. Journal of Bacteriology. 15 juin 1998;180(12):3107–13.

59. Brown DG, Swanson JK, Allen C. Two Host-Induced Ralstonia solanacearum Genes, acrA and dinF, Encode Multidrug Efflux Pumps and Contribute to Bacterial Wilt Virulence. Applied and Environmental Microbiology. mai 2007;73(9):2777–86.

60. Pletzer D, Weingart H. Characterization and regulation of the Resistance-Nodulation-Cell Division-type multidrug efflux pumps MdtABC and MdtUVW from the fire blight pathogen Erwinia amylovora. BMC Microbiology. 11 juill 2014;14(1):185.

61. Vargas P, Felipe A, Michán C, Gallegos MT. Induction of Pseudomonas syringae pv. tomato DC3000 MexAB-OprM Multidrug Efflux Pump by Flavonoids Is Mediated by the Repressor PmeR. MPMI. oct 2011;24(10):1207–19.

62. Badri DV, Vivanco JM. Regulation and function of root exudates. Plant, Cell & Environment. 2009;32(6):666–81.

63. Tiziani R, Miras-Moreno B, Malacrinò A, Vescio R, Lucini L, Mimmo T, et al. Drought, heat, and their combination impact the root exudation patterns and rhizosphere microbiome in maize roots. Environmental and Experimental Botany. 1 nov 2022;203:105071.

64. Zhalnina K, Louie KB, Hao Z, Mansoori N, da Rocha UN, Shi S, et al. Dynamic root exudate chemistry and microbial substrate preferences drive patterns in rhizosphere microbial community assembly. Nat Microbiol. avr 2018;3(4):470–80.

65. Ravirala RS, Barabote RD, Wheeler DM, Reverchon S, Tatum O, Malouf J, et al. Efflux Pump Gene Expression in Erwinia chrysanthemi Is Induced by Exposure to Phenolic Acids. MPMI. mars 2007;20(3):313–20.

66. Jordana-Lluch E, Barceló IM, Escobar-Salom M, Estévez MA, Zamorano L, Gómez-Zorrilla S, et al. The balance between antibiotic resistance and fitness/virulence in Pseudomonas aeruginosa: an update on basic knowledge and fundamental research. Front Microbiol [Internet]. 28 sept 2023 [cité 29 mai 2024];14. Disponible sur: https://www.frontiersin.org/journals/microbiology/articles/10.3389/fmicb.2023.1270999/full

